# Distinct molecular targets for macrocyclic lactone and isoxazoline insecticides in the human louse: new prospects for the treatment of pediculosis

**DOI:** 10.1101/2020.08.06.239400

**Authors:** Nicolas Lamassiaude, Berthine Toubate, Cédric Neveu, Pierre Charnet, Catherine Dupuy, Françoise Debierre-Grockiego, Isabelle Dimier-Poisson, Claude L. Charvet

**Author notes:** Co-corresponding authors (CLC), Institut national de recherche pour l’agriculture, l’alimentation et l’environnement, UMR1282 - Infectiologie et Santé Publique, Centre de Recherche Val de Loire, 37380 Nouzilly, France, (IDP), Université de Tours, UMR1282 - Infectiologie et Santé Publique, 37000 Tours, France.

## Abstract

**Background:** Control of infestation by cosmopolitan lice (*Pediculus humanus*) is increasingly difficult due to the transmission of resistant parasites to pediculicides. However, since the targets for pediculicides have no been identified in human lice so far, their mechanism of action remain largely unknown. The macrocyclic lactone, ivermectin is active against a broad range of insects including human lice. Isoxazolines are a recent new chemical class targeting the γ-aminobutyric acid (GABA) receptor made of the <u>r</u>esistant to <u>d</u>ie<u>l</u>drin (RDL) subunit and exhibiting a strong insecticidal potential. Here, we addressed the pediculicidal potential of isoxazolines and deciphered the molecular targets of ivermectin and the novel ectoparasiticide lotilaner in the human body louse species *Pediculus humanus humanus*.

**Methods and findings:** Using toxicity bioassays, we showed that fipronil, ivermectin and lotilaner are efficient pediculicides on adult lice. The RDL (Phh-RDL) and glutamate-gated chloride channel subunits (Phh-GluCl) were cloned and characterized by two-electrode voltage clamp electrophysiology in *Xenopus laevis* oocytes. Phh-RDL and Phh-GluCl formed functional homomeric receptors respectively gated by GABA and L-glutamate with EC_50_ values of 6.4 µM and 9.3 µM. Importantly, ivermectin displayed a super agonist action on Phh-Glucl whereas Phh-RDL receptors were weakly affected. Reversally, lotilaner strongly inhibited the GABA-evoked currents in Phh-RDL with an IC_50_ value 0.5 µM, whereas it had no effect on Phh-GluCl.

**Conclusions:** We report here for the first time the pediculicidal potential of isoxazolines and reveal the distinct mode of action of ivermectin and lotilaner on GluCl and RDL channels from human lice. These results emphasize the clear repurposing future of the isoxazoline drug class to provide new pediculicidal agents to tackle resistant-louse infestations in humans.

**Autorship summary:** *Why was this study done?:* Human cosmopolitan lice are responsible for pediculosis, which represent a significant public health concern. Resistant lice against insecticides and lack of safety of the treatments for human and environment is a growing issue worldwide. Here we investigated the efficacy on lice of the classical macrocyclic lactone drug, ivermectin, and the novel isoxazoline drug, lotilaner. This study was done to decipher their mode of action at the molecular and funtional levels in order to propose new strategies to control lice infestation.

*What did the researchers do and find?:* Our bioassay results indicate that ivermectin and lotilaner were potent at killing human adult lice, with lotilaner showing a higher efficacy than ivermectin. Furthermore, we identified and pharmacologically characterized the first glutamate- and GABA-gated chloride channels ever described in human lice yet. Mechanistically, our molecular biology and electrophysiology findings demonstrate that ivermectin acted preferentially at glutamate channels while lotilaner specifically targeted GABA channels.

*What do these findings mean?:* These results provide new insights in the understanding of the insecticide mode of action and highlight the isoxazolines as a new drug-repurposing opportunity for pest control strategies.

## Introduction

Lice are spread throughout the world in both low- and high-income countries, with heterogenous prevalence depending on the geographical area [1, 2]. Pediculosis prevailing among school age children and homeless persons represents major economic, social but also public health concerns [3-8]. Human lice include the body louse *Pediculus humanus humanus* and the head louse *Pediculus humanus capitis* (order: *Phthiraptera*: Pediculidae). Both species are obligate blood feeding ectoparasites, living in clothes and in the scalp area, respectively [9]. Despite morphological, physiological and ecological differences, genetic studies showed that head and body lice have almost the same genomic and transcriptomic contents [9-11]. Furthermore, body lice are vectors of the pathogenic bacteria *Bartonella quintana, Borrelia recurrentis* and *Rickettsia prowazekii*, responsible for trench fever, louse-borne relapsing fever and epidemic typhus, respectively [4]. The control of lice infestation remains increasingly difficult due to the mode of transmission and the relative unreliable efficicacy of the available treatments [12]. Even though silicon-based suffocating products are widely used nowadays, their efficacy largely depends on the respect of their application and is limited against the nits [13]. Chemical insecticides such as organochloride (lindane), organophosphates (malathion), carbamates (carbaryl) and pyrethroids (pyrethrins and synthetic permethrin), which are the first line treatments recommended by the Centers for Disease Control and Prevention and by the American Academy of Pediatrics, have been widely used to control lice [14, 15]. They have been proposed to inhibit γ-aminobutyric acid (GABA)-gated chloride channels, acetylcholinesterase and voltage-gated sodium channels, respectively, leading to the paralysis of the parasite. However, louse resistance to these products has been widely reported [16-19] as well as neurotoxic effects in children and toxicity on the environment [20-25]. Because macrocyclic lactones (MLs) are broad-spectrum neuroactive insecticides, acaricides and nematicides used to treat and prevent endo- and ectoparasites of humans and animals [26], ivermectin (a derivate of the ML avermectins) has emerged as a promising pediculicide and quickly became the recommended molecule for the control of body lice [27-29]. In invertebrates, ivermectin inhibits glutamate-gated chloride channels (GluCls) leading to a reduction in motor activity, paralysis and death. However, the molecular basis for the action of ivermectin in louse is unknown. Recently, treatment failures of ivermectin against lice were observed [30, 31]. In that respect, there is an urgent need to identify new druggable targets and develop new pediculicidal compounds.

Cys-loop ligand-gated ion channels (LGICs) are the major pharmacological targets of insecticides [32]. Among LGICs, the GluCls are found at inhibitory synapses of the nervous system exclusively in invertebrates [33], while γ-aminobutyric acid (GABA) is the main inhibitory neurotransmitter in nerves and muscles in both vertebrates and invertebrates. GluCls and GABA receptors are constituted by the oligomerization of five identical or different subunits around a central pore [34]. In arthropods, the finding that the insecticidal activity of avermectins is mainly mediated by the GluCls was first achieved in the model insect *Drosophila melanogaster* [35]. For the GABA channels (GABACls), the screening for mutant flies that survive exposure to the insecticides dieldrin and fipronil, allowed the identification of the *resistant to <u>d</u>ieldrin* gene or *rdl* [36]. When heterologously expressed in *Xenopus laevis* oocytes, the GluCl and RDL subunits formed functional homomeric glutamate-gated and GABA-gated chloride channels, respectively [35, 37]. GluCl and RDL subunits were subsequently described in many other insect species although few of them were functionally assayed *in vitro*. Mutations in the GluCl subunit confers resistance to avermectins, whereas resistance to dieldrin, fipronil and picrotoxinin was shown to be associated to mutations in the RDL subunit using functional *in vitro* and *in vivo* experiments [32].

Interestingly, new synthetic molecules from the isoxazoline chemical class were recently proven to be effective ectoparasiticides against fleas and ticks [38]. The isoxazoline insecticides include fluralaner, afoxolaner, sarolaner and lotilaner [39-42]. They were shown to act as potent non-competitive antagonists of GABACls from *D. melanogaster* as well as the cat flea *Ctenocephalides felis* and the livestock tick *Rhipicephalus microplus* by molecular experiments and voltage-clamp electrophysiology. In addition, these compounds were efficient at blocking GABA-elicited currents from dieldrin- and fipronil-resistant channels. Recent studies on the actions of these compounds on cloned GABACls have accounted for their higher selectivity for insect over vertebrate receptors [42, 43]. Compounds from the isoxazoline group are now considered as the next generation of broad-spectrum insecticides effective against veterinary, agricultural, sanitary and stored product pests from eleven arthropod orders [44]. However, the pediculicidal activity of isoxazolines, the identification of the target ion channels in lice and their potential use for human health have not been investigated so far.

The objectives of our study were to decipher the molecular targets of ivermectin and lotilaner in the human body louse species *Pediculus humanus humanus* (Phh). In the present paper, we describe in the human body louse: the *in vivo* pediculicidal activity, the target identification and *in vitro* mechanism of action of the old phenylpyrazole insecticide, fipronil, the recent ML insecticide, ivermectin, and the novel isoxazoline, lotilaner. For this, we identified the human body louse orthologues of the GluCl and RDL subunit genes and subsequently achieved their respective functional characterizations in *Xenopus* oocytes using two-electrode voltage-clamp. We found that the Phh-GluCl and Phh-RDL subunits give rise to glutamate-gated and GABA-gated homomeric channels having distinct pharmacological properties. Hence, we provide mechanistic insights for a distinct mode of action of ivermectin and lotilaner. In the context of drug-resistance, this study highlights the pediculicidal potential of lotilaner in a drug-repurposing attempt of new strategies to control human louse infestations.

## Results

### *In vivo* toxicity tests of ivermectin and lotilaner on lice and nits

In order to determine the efficacy of selected insecticides, we first performed insecticide exposure tests on laboratory-reared human adult body lice and nits (**Fig. 1A**). Briefly, lice were incubated with ivermectin, fipronil, picrotoxin and lotilaner at different concentrations. The effects of the compounds on louse survival and immobilization (knockdown) were subsequently analyzed at four timepoints (1, 2, 3, and 24 hours). After 24h, there was 4.5% mortality in the groups of lice treated with 1% DMSO that was used to dissolve the different compounds (**Fig. 1B**). Similarly, 100 µM picrotoxin was not efficient to kill the lice after 24h whereas 91% of the lice were killed or immobilized with exposure to 100 µM fipronil In our experiments, fipronil was used as a positive control and this result corroborated previous bioassays in the literature [45]. As expected, ivermectin was more efficient than fipronil, since the death rate reached 30%, 3h after application of 100 µM, and 100% after 24h application of both 10 and 100 µM (**Fig. 1B)**. These findings confirmed the insecticide susceptible status of the laboratory-reared colony used here. Strikingly, after application of 100 µM lotilaner, the number of living lice declined rapidly. Not only the application of 1 µM lotilaner killed 14% of lice at 24 h but 10 and 100 µM lotilaner led to the death of all lice 3h after exposition (**Fig. 1B**). Interestingly, when compared with other drugs, 10 µM lotilaner was more efficient than 10 µM ivermectin and 100 µM fipronil. Next, we analyzed the effect of 100 µM ivermectin or lotilaner on the nit hatching between six and nine days after drug application. The rate of nits hatched did not differ between the ivermectin- and the lotilaner-treated nits and the negative controls (water and 1% DMSO) thus revealing no effect of the two products on the nits (**Fig. 1C**). Altogether, these results confirm that fipronil and ivermectin are efficient at killing human adult body lice and provide the first *in vivo* evidence of a pediculicidal activity for lotilaner. These data prompted us to investigate the targets of ivermectin and lotilaner at the molecular level.

**Fig. 1:**
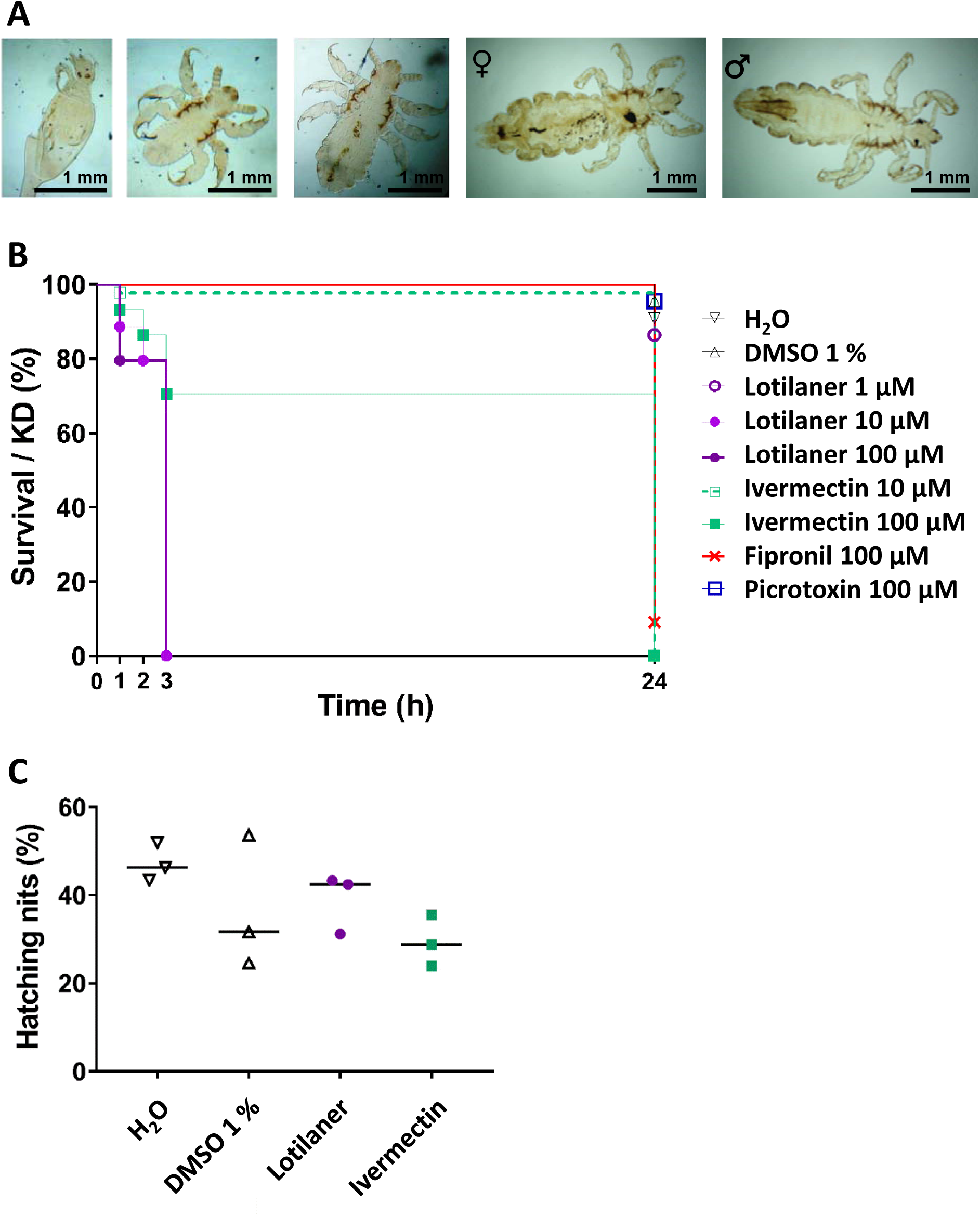
Toxicity tests on lice *in vivo*. **A**. Photographs of the human body lice from the laboratory-reared colony used in this study. Different life stages are represented from the left to the right: larvae in stage 1 during hatching, larvae in stage 2, larvae in stage 3, female adult, male adult. **B. A**dult lice (n = 44 per condition) were treated with different concentrations of lotilaner, ivermectin, fipronil or picrotoxin and survival/immobilization (knockdown) rate was determined throughout 24 hours. Water and 1 % DMSO were used as negative controls. The results are representative of three independent experiments. **C**. Nits were treated with 100µM of lotilaner or ivermectin and hatching was determined throughout several days. Water and 1 % DMSO were used as negative controls. The results are expressed as percentage of hatching from three independent experiments with n = 50 to 190 nits.

### GluCl and RDL subunits are conserved in *P. humanus humanus*

Macrocyclic lactones and isoxazolines were previously shown to act on arthropod glutamate- and GABA-gated channels. Therefore, we decided to identify and clone the respective *P. humanus humanus* GluCl and RDL subunits through a candidate gene approach. Searches for homologues of *A. mellifera* and *D. melanogaster* GluCl and RDL receptor subunits in *P. humanus* genomic/transcriptomic databanks, allowed the identification of two independent coding sequences for *Phh-GluCl* and for *Phh-rdl* for which the respective complete coding sequences were obtained by RACE PCR experiments. Both louse GluCl and RDL displayed the characteristic features of cys-loop LGIC subunits, including a signal peptide, an extracellular N-terminal domain containing the ligand binding sites, a disulfide bond formed by two cysteines that are 13 amino acid residues apart (cys-loop domain), four transmembrane domains (TM1-TM4) and a large intracellular loop between TM3 and TM4 that is highly variable (**Fig. 2**). Of importance is the sequence analysis of the pore selectivity filter located in the TM2 supporting the prediction with the signature of a chloride channel. Indeed, Phh-GluCl and Phh-RDL both contain the PAR motif centered at the 0’ position in the TM2 and conserved in other species (universal LGIC TM2 numbering system). A proline at position −2’, followed by a small amino acid at position −1’ like an alanine, an arginine at the 0’ residue and a threonine at position 13’ are the hallmarks of chloride–permeable LGICs [46].

**Fig. 2:**
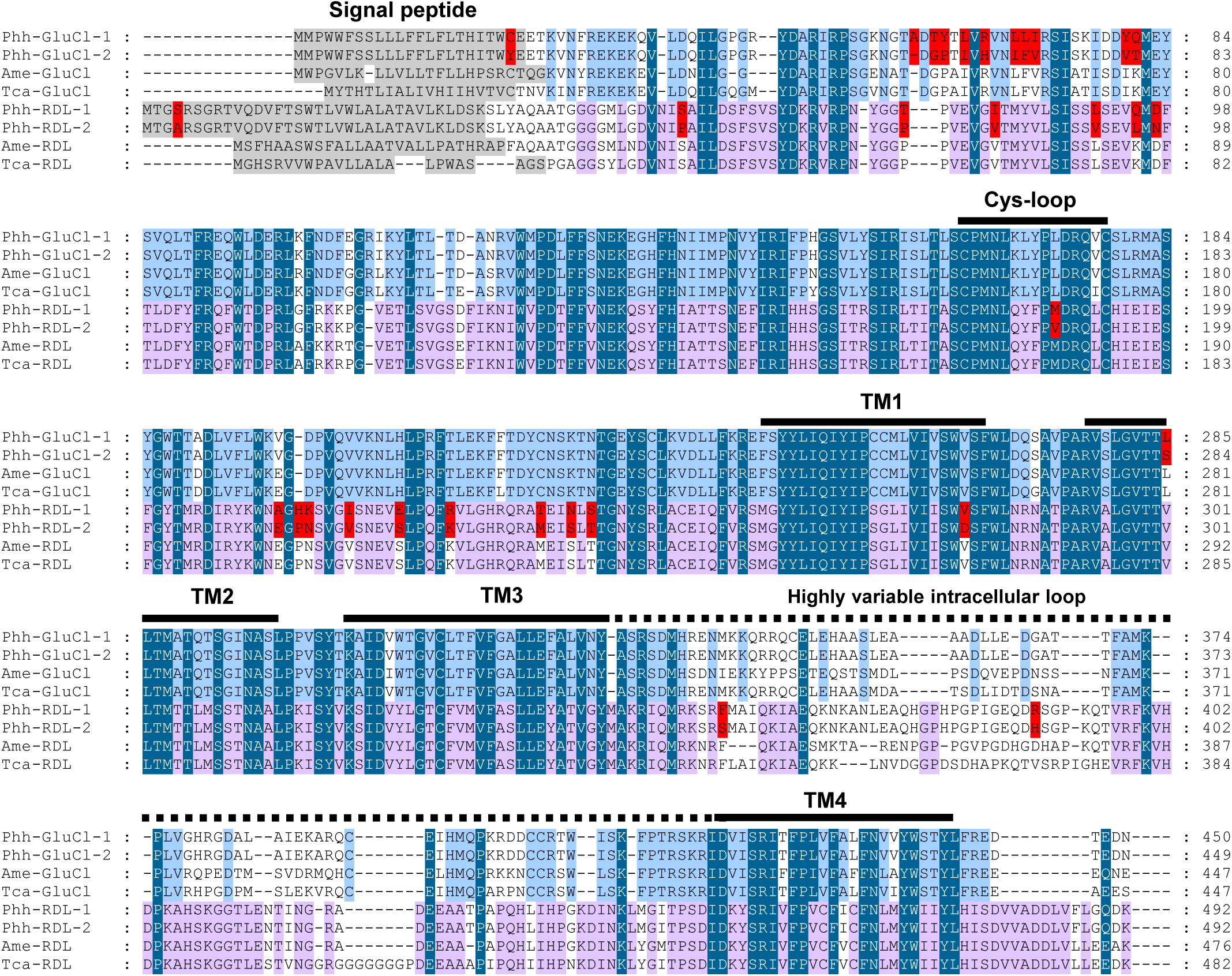
Amino acid alignments of GluCl and RDL subunit sequences of *Pediculus humanus humanus, Apis mellifera* and *Tribolium castaneum*. Predicted signal peptides in the N-terminal region are highlighted in grey. Amino acid differences between Phh-GluCl-1 and Phh-GluCl-2 or between Phh-RDL-1 and Phh-RDL-2 sequences of *P. humanus humanus* are highlighted in red. Cys-loop domain, predicted transmembrane domains TM1 to TM4 and the highly variable intracellular loop are indicated by the bars. Amino acids conserved between all the sequences are highlighted in dark blue. Amino-acids conserved between GluCl sequences are highlighted in light blue. Amino acids conserved between RDL sequences are highlighted in pink.

The full-length coding sequences for the GluCl subunits included ORFs of 1353 and 1350 pb encoding proteins of 450 and 449 amino acids that were named Phh-GluCl-1 (MT321070) and Phh-GluCl-2 (MT321071), respectively. The alignement of *Phh-GluCl-1* and *Phh-GluCl-2* deduced amino-acid sequences is provided in **Fig. 2**. The two sequences are highly conserved with 97% identity and 98% similarity and only differ by eleven amino acid substitutions and an alanine insertion in the N-terminal part at position 59 in Phh-GluCl-1. (**Fig. 2** and **S1 Fig**.). Noteworthy, all the amino acids involved in the glutamate binding were conserved.

For RDL subunits, both full-length coding sequences are 1467 nucleotide long encoding 489 amino acids. Phh-RDL-1 (MT321072) and Phh-RDL-2 (MT321073) share 96% identity and 98% similarity, only differing by 20 amino acid substitutions including 17 amino acid changes located in the extracellular N-terminal region between the signal peptide and the TM1 (**Fig. 2** and **S2 Fig**.). Notably, none of the residues involved in the binding of GABA were subjected to substitutions.

A distance tree analysis of *P. humanus humanus* GluCl and RDL sequences including *D. melanogaster*, the honey bee *Apis mellifera*, the house fly *Musca domestica, R. microplus* and the red flour beetle *Tribolium castaneum* confirmed the orthologous relationships of the louse subunit sequences with their respective counterparts (**Fig. 3**). GluCl amino acid sequences revealed that Phh-GluCls were found to be highly conserved, sharing 78%, 84%, 81% and 70% identities with the respective orthologues from *A. mellifera, T. castaneum, D. melanogaster* and *R. microplus*. Phh-RDL-1 and 2 shared 84%, 81%, 67% and 54% identities with their orthologues from *A. mellifera, T. castaneum, D. melanogaster* and *R. microplus*, respectively. These observations are in accordance with the relative position of Phthiraptera and Hymenoptera insect subfamilies and Acarids [47] and were further confirmed by supplementary analysis including other members from the GABA and GluCl groups from additional arthropod species (**S1 Fig**. and **S2 Fig**.). In the phylogeny of GABA subunits, Phh-RDL-1/-2 were found to clearly cluster distinctly from the LCCH3, GRD and 8916 subunits, showing that each species has inherited an ancestral gene. Altogether, the phylogenetic observations support that we cloned the *P. humanus humanus* GluCl and RDL subunit orthologues allowing their subsequent assessment at the functional level.

**Fig. 3:**
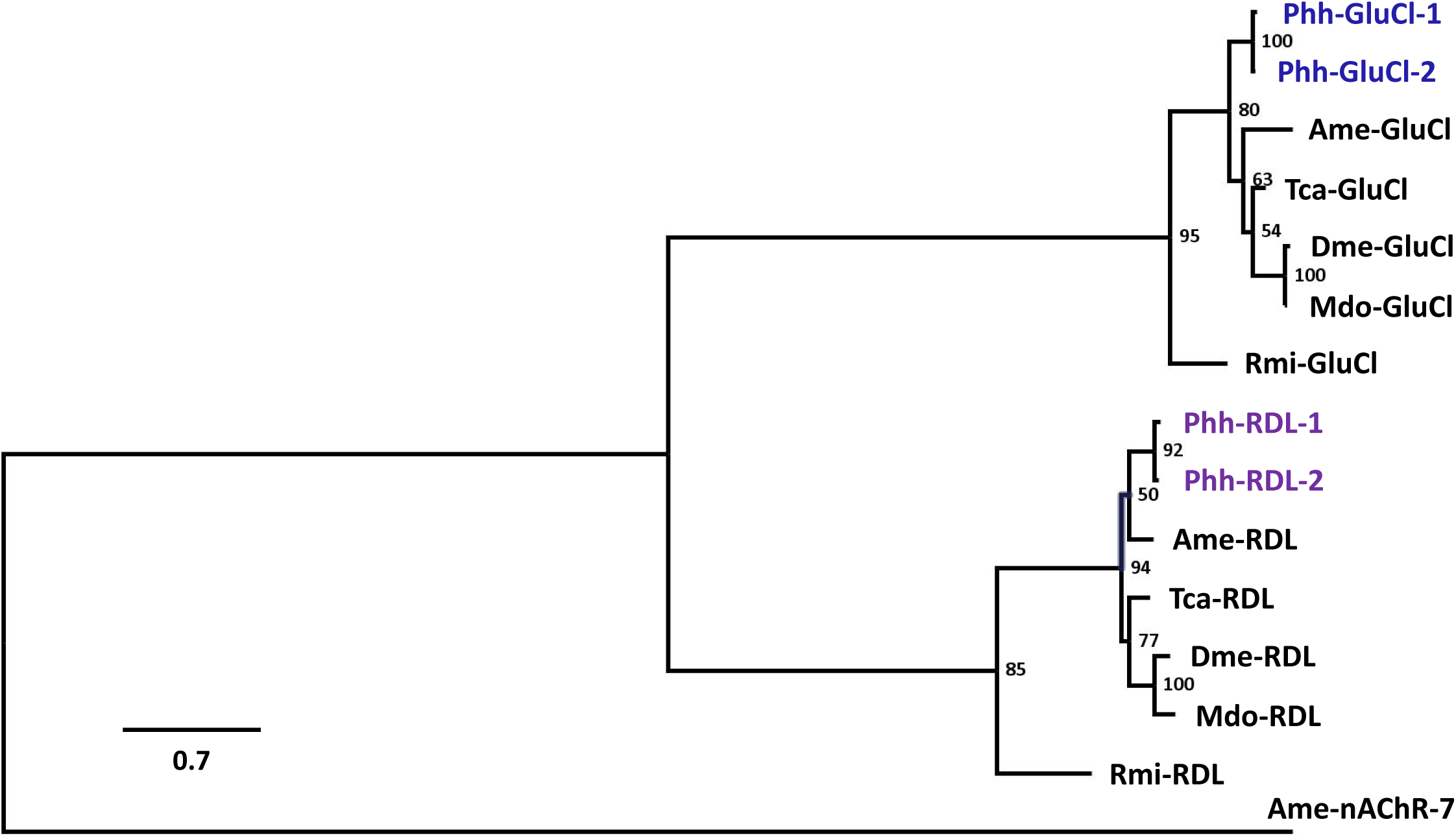
Distance tree (BioNJ, Poisson) of GluCl and RDL protein sequences from insects and acarians. The three letter prefixes in gene names Phh, Ame, Tca, Dme, Mdo and Rmi refer to the species *Pediculus humanus humanus, Apis mellifera, Tribolium castaneum, Drosophila melanogaster, Musca domestica* and *Rhipicephalus microplus*, respectively. Branch lengths are proportional to the number of substitutions per site. Scale bar represents the number of substitutions per site. The bootstrap values are indicated next to each branch. Accession numbers for sequences used in the phylogenetic analysis are provided in the Methods section. Sequences of *P. humanus humanus* are highlighted in blue for GluCls and purple for RDLs. The *A. mellifera* alpha7 nAChR subunit sequence was used as an outgroup.

### *P. humanus humanus* GluCl and RDL subunits form functional homomeric channels in *Xenopus* oocytes

In previous studies, arthropod GluCl and RDL subunits were shown to reconstitute functional channels when expressed in *Xenopus* oocytes [32]. Hence, we hypothesized that the ortholog subunits from *P. humanus humanus* could form functional receptors. Following the cloning of two full-length coding sequences for Phh-GluCl and Phh-RDL subunits, their respective cRNAs were microinjected in *X. laevis* oocytes. To investigate the receptor functionality, responses to agonists were assessed by two-electrode voltage-clamp (TEVC) electrophysiology. In sharp contrast with Phh-GluCl-1 and Phh-RDL-1 subunits (**Fig. 4**), injections of *Phh-GluCl-2* (n=11) and *Phh-RDL-2* (n=12) cRNAs never led to recordings of glutamate- and GABA-evoked currents, respectively, indicating that these subunits did not form functional receptors (**S3 Fig**.). Thus, Phh-GluCl-1 and Phh-RDL-1 will be henceforth refered to as Phh-GluCl and Phh-RDL in the article. When glutamate was perfused in the recording chamber, we could record fast inward currents in the µA range in oocytes injected with *Phh-GluCl* (**Fig. 4A)**. As expected, this receptor desensitized extremely rapidly upon prolonged glutamate exposure. Then, we obtained the concentration-response curve by challenging the oocytes with glutamate concentrations ranging from 1 to 1000 µM. With current amplitudes normalized to the maximal response to 1000 µM, we found that the glutamate EC_50_ value of Phh-GluCl was 9.3 +/- 1.3 µM with a Hill coefficient of 1.9 +/- 0.5 (n=5) suggesting the presence of more than one glutamate binding site (**Fig. 4A-B)**. When *Phh-RDL* cRNAs were injected in oocytes, the application of 100 µM GABA resulted in robust currents with maximum amplitudes in the µA range (**Fig. 4C**). The GABA concentration-response curve was characterized by an EC_50_ of 6.4 +/- 0.4 µM (n=6) with a Hill coefficient of 2.1 +/- 0.2 (n=6) suggesting the binding of more than two GABA molecules (**Fig. 4C-D**). As previously shown in the literature, RDL and GluCls are permeable to chloride anions [35, 37, 48-51]. Here, the TM2 amino acid sequences of Phh-RDL and Phh-GluCl are respectively 100% identical with RDL and GluCls from other insect species such as *A. mellifera, D. melanogaster, T. castaneum* (**Fig. 2**), suggesting they are anionic receptors. In order to investigate the ion selectivity, we applied voltage-ramps to oocytes expressing Phh-RDL and measured the reversal potentials for GABA-sensitive currents in recording solution with varying concentrations of either chloride or sodium. Then, the reversal potentials were plotted against the chloride or sodium concentration (**Fig. 4E**). In the standard recording solution containing 100 mM NaCl, GABA-induced currents reversed at −22.5 +/- 0.8 mV (n=10) (**Fig. 4E**). When the extracellular chloride concentration decreased from 100 to 0 mM (replacement of chloride by sodium acetate), Phh-RDL exhibited a 36.4 ± 1.2 mV shift upon a 10-fold change in the chloride concentration (n=4). In contrast, decreasing the extracellular sodium concentration from 100 to 0 mM (replaced by tetraethylammonium chloride) did not significantly shift the reversal potential (2.3 ± 0.8 mV shift per decade, n=4). These results indicate that Phh-RDL forms a chloride permeable GABA-gated channel that does not permeate sodium ions, consistently with the pore selectivity filter sequence (**Fig. 2**) and RDL channels from other species [51, 52]. In summary, these findings demonstrate that two novel functional homomeric receptors from the human body louse, Phh-GluCl and Phh-RDL, with respectively high affinity to glutamate or GABA were reconstituted in *Xenopus* oocytes.

**Fig. 4:**
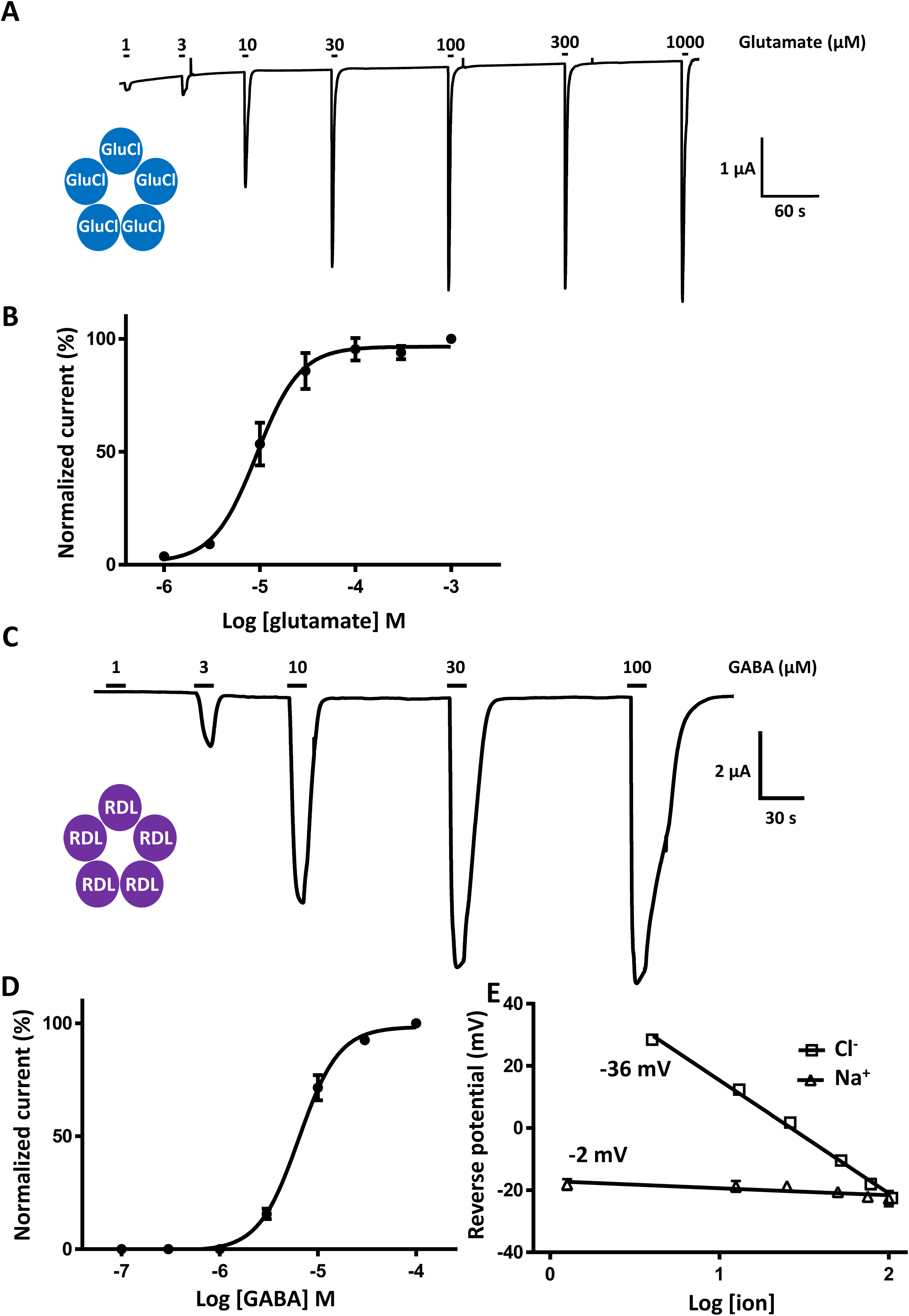
Functional expression of Phh-GluCl and Phh-RDL channels in *X. laevis* oocytes. **A**. Electrophysiological recording traces of oocytes injected with Phh-GluCl cRNA in response to the application of 1 to 1000 µM of glutamate. Oocytes were clamped at −80 mV. Application times are indicated by the bar. **B**. Glutamate concentration-response curve for Phh-GluCl (mean +/- SEM, n = 5 oocytes). Data were normalized to the maximal effect of glutamate. **C**. Electrophysiological recording traces of oocytes injected with Phh-RDL cRNA in response to the application of 1 to 100 µM of GABA. Oocytes were clamped at −60 mV. Application times are indicated by the bars. **D**. GABA concentration response curve for Phh-RDL (mean +/- SEM, n = 6 oocytes). Data were normalized to the maximal effect of GABA. **E**. Reverse potential-ion concentration relationships of oocytes expressing Phh-RDL obtained from voltage-ramps in presence of GABA and using increasing concentrations of either Cl- or Na+ to analyze the selectivity of this channel.

### Differential effects of ivermectin and lotilaner on *P. humanus humanus* GluCl and RDL channels

Arthropod GluCl and RDL channels are well-known targets for insecticides [32]. In order to investigate their respective insecticides sensitivities, Phh-GluCl and Phh-RDL were challenged with ivermectin, lotilaner, fipronil and picrotoxin. Recordings from oocytes expressing Phh-GluCl revealed that ivermectin is a potent agonist of this receptor. A slow activation of the receptor was observed with application of ivermectin as low as 10 nM and large µA currents were recorded in response to 1 µM ivermectin with an average of 86.5 +/- 8.2% (n=6) of the 1 mM glutamate-elicited current (**Fig. 5A-B**). As expected, the effect of ivermectin was not reversible. When applied on Phh-GluCl expressing oocytes, 1 µM lotilaner did not elicit any current and mediated no inhibitory effect on glutamate-evoked currents (**Fig. 5C**). We subsequently tested the activity of selected insecticides on the Phh-RDL channel. As anticipated from the litterature, ivermectin, lotilaner, picrotoxin and fipronil had no agonist effect (**Fig. 6** and **Fig. S4**). Then, we investigated the potential antagonistic effects of the selected insecticides on GABA recording traces. **Figures 6A-D** and **S4** illustrate results from application of 100 µM GABA before, during and after the addition of either 10 µM or 1 µM of insecticides, respectively. When applied at 1 µM fipronil, picrotoxin and lotilaner partially antagonized GABA-evoked currents while 1 µM ivermectin produced an almost insignificant effect (**Fig. S4**). Rank order potency series for each of the Phh-RDL antagonists tested at a concentration of 1 μM was as follows: lotilaner > picrotoxin ≈ fipronil > ivermectin. When perfused at 10 µM, the selected insecticides decreased significantly the GABA-evoked current amplitude (**Fig. 6**). Fipronil and picrotoxin blocked 92 +/- 4 % (n = 6) and 82 +/- 5 % (n = 5) of the GABA response on Phh-RDL respectively (**Fig. 6A-B** and **6E-F**). Surprinsingly, ivermectin acted as a weak inhibitor of Phh-RDL with a partial effect of only 31 +/- 5 % (n=9) (**Fig. 6C** and **6E**). More importantly, GABA responses were completely and irreversibly blocked with lotilaner as low as 3 µM suggesting that Phh-RDL is more sensitive to lotilaner than to the ancient insecticides fipronil or picrotoxin (**Fig. 6D-F**). To characterize this effect, antagonist concentration-response relationships were perfomed (**Fig. 6F**). Fipronil and picrotoxin antagonized Phh-RDL in a concentration-dependent manner with IC_50_ values of 1.4 +/- 0.3 µM (n = 6 oocytes) and 2.8 +/- 1.4 µM (n = 5 oocytes), respectively. Similarly, lotilaner resulted in the most potent dose-dependent inhibition of the GABA-mediated currents with an IC_50_ value of 0.5 +/- 0.3 µM (n = 6 oocytes). Altogether, these results demonstrate that *P. humanus humanus* GluCl and RDL characterized in *Xenopus* oocytes display unique and distinct pharmacologies regarding insecticides. Ivermectin acted as a strong agonist Phh-GluCl and a weak antagonist of Phh-RDL, while the novel anti-ticks and fleas lotilaner was a potent and selective blocker of RDL of human lice.

**Fig. 5:**
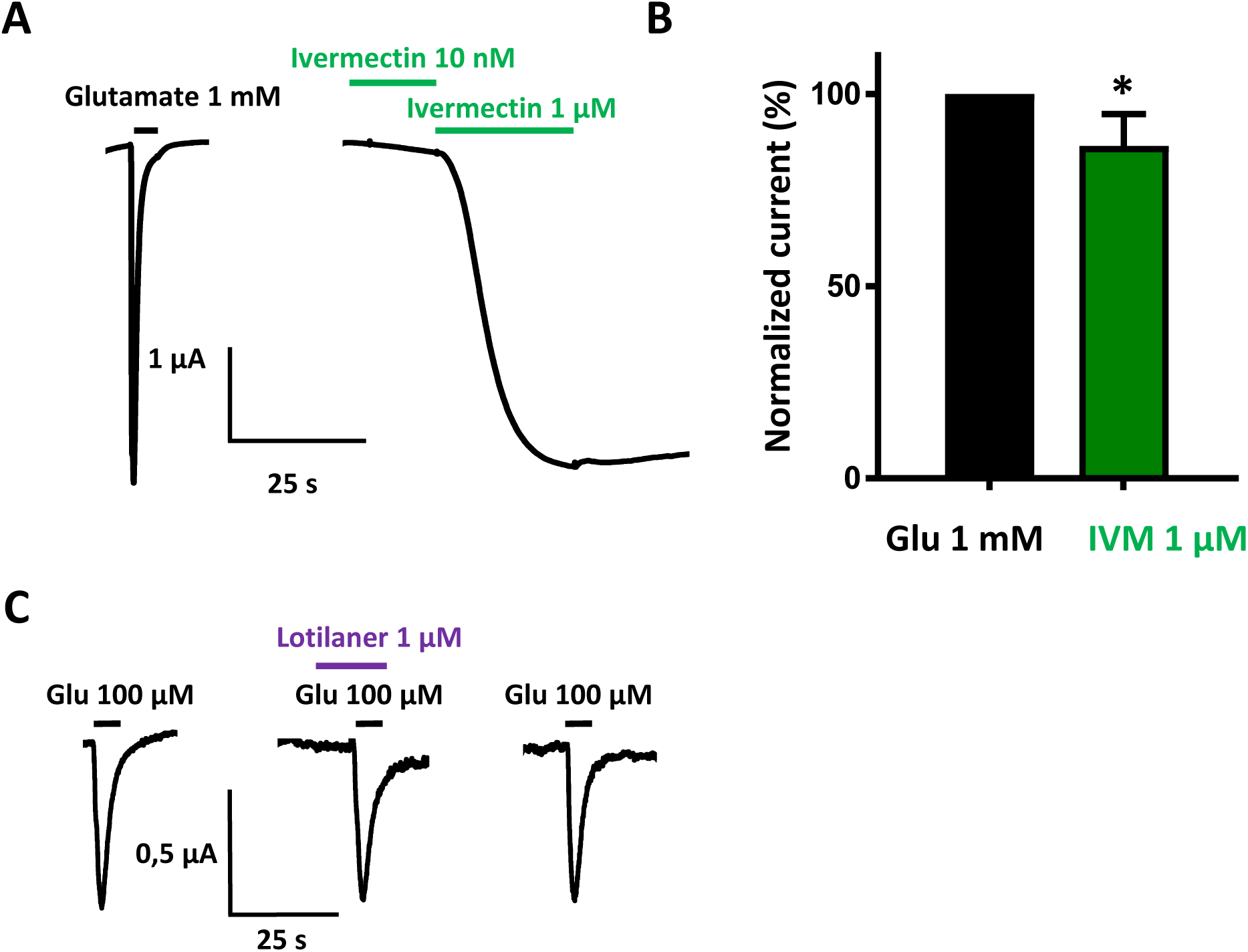
Effects of ivermectin and lotilaner on Phh-GluCl receptor in *X. laevis* oocytes. **A**. Representative recording traces from a single oocyte expressing Phh-GluCl challenged with 1 mM glutamate and 10 nM ivermectin followed by 1 µM ivermectin. The bars indicate the time period of agonist application. **B**. Bar chart (mean +/- SEM, n = 6 oocytes) of the response elicited by 1 µM ivermectin (IVM) normalized to and compared with the maximal response to 1 mM glutamate (Glu). *, *p* < 0.05; Student’s t-test **C**. Representative recording traces from a single oocyte perfused with 100 µM glutamate (Glu) alone (left), with 1 µM lotilaner prior to co-application with 100 µM Glu (middle) and after wash out (right).

**Fig. 6:**
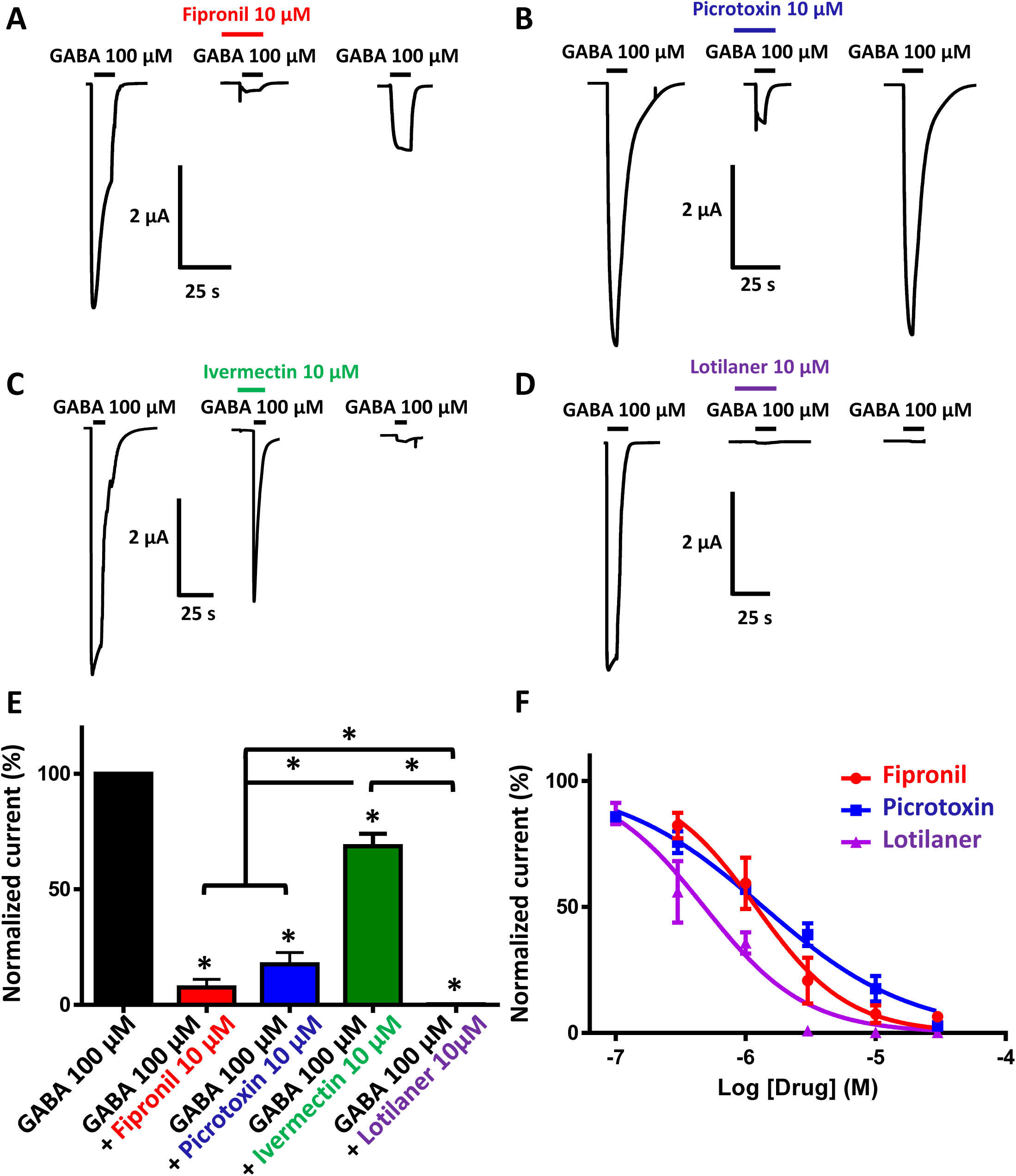
Antagonistic effects of fipronil, picrotoxin, ivermectin and lotilaner on GABA-elicited currents in *X. laevis* oocytes expressing Phh-RDL receptor. **A-D**. Representative current traces evoked by 100 µM GABA with or without co-application of 10 µM fipronil, picrotoxin, ivermectin and lotilaner. Application times are indicated by the bars. **E**. Bar chart of the normalized current responses for 100 µM GABA alone and in the presence of 10 µM fipronil, picrotoxin, ivermectin and lotilaner on Phh-RDL (mean +/- SEM, n = 3-9 oocytes). Currents have been normalized to and compared with 100 µM GABA-elicited currents. *, *p* < 0.05; One-way ANOVA with Tukey’s Multiple Comparisons Test. **F**. Concentration-inhibition curves for Phh-RDL for fipronil (red circles, n = 6-7), picrotoxin (blue squares, n = 5-8) and lotilaner (purple triangles, n = 3-14). The responses are all normalized to the response to 100 µM GABA (mean +/- SEM, n = 3-14 oocytes).

## Discussion

### First functional characterization of LGICs from human lice

In the present study, we report the first functional expression and pharmacological characterization of any human louse LGICs. Indeed, despite the social and public health impacts of human lice and the growing issue of resistance to pediculicides and since the complete sequencing of the body louse genomes, there was to our knowledge no functional characterization of any LGICs from lice so far [10, 29]. Because glutamate and GABA are important neurotransmitters in the insect central nervous system and their receptors are targets for insecticides, we focused our attention on glutamate- and GABA-sensitive receptors. GluCl and RDL channel subtypes of LGICs have been extensively characterized in many arthropod species starting with the model fruit fly *D. melanogaster* and ranging from agricultural and sanitary pests to animal ectoparasites and disease vectors but also beneficial insects like the honey bee [32, 33]. Here, we show that *P. humanus humanus* GluCl and RDL subunits gave rise to homomeric channels when functionally expressed in *Xenopus* oocytes. This experimental expression system provided a unique means to assess the pharmacological and biophysical properties of these receptors. Even though two putative genes encoding respectively a GluCl and a GABA receptor have been identified in the genomic database, the molecular cloning of complete cDNAs revealed at least two potential full-length transcripts for both candidates (i.e. Phh-GluCl-1, Phh-GluCl-2 and Phh-RDL-1, Phh-RDL-2). These transcripts could either be alternative spliced isoforms or result from RNA A-to-I editing, a mechanism of RNA edition described in lice [53]. In insects, alternative splicing and RNA A-to-I editing are known to increased diversity of LGICs subunits [54-57]. Undoubtedly, further molecular investigations at the DNA level would be required to confirm this hypothesis. Here, we report that both Phh-GluCl-1 and Phh-RDL-1 can be robustly expressed in *Xenopus* oocytes, whereas neither Phh-GluCl-2 nor Phh-RDL-2 subunits formed functional receptors in our conditions. Interestingly, Phh-GluCl had one of the highest affinity for glutamate compared to other GluCls of arthropod species, the reported EC_50_ of which varied between 6.89 and 384.2 µM [35, 43, 48-50, 58-62]. Likewise, Phh-RDL had one of the lowest EC_50_ for GABA among described insect RDL receptors showing EC_50_ varying between 5.43 +/- 0.02 µM in the brown planthopper *Nilaparvata lugens* [63] and 445 +/- 47 µM in the carmine spider mite *Tetranychus cinnabarinus* [61] with the exception of the RDL-2 subunit (2.4 +/- 0.3 µM) from the honey bee parasite mite *Varroa destructor* [51]. Similarly, spliced variant transcripts of GluCl and RDL subunits were found to give rise to functional channels in several insect species, thus increasing their receptor diversity [33, 56, 57, 60, 62, 64-66]. Here, we identified and characterized the first GluCl and GABACl receptors of human lice. This study opens the way for the characterization of other receptor subtypes which can also be relevant target.

### New insights into the mode of action of ivermectin on lice

Despite the massive use of pediculicidal compounds during the past decades, there was to our knowledge no functional characterization of their molecular targets in lice. Macrocyclic lactones are widely used in both veterinary and human medicine, with ivermectin being typically the recommended molecule for the control of body lice [27-29]. This broad-spectrum drug is highly efficient against a wide range of pathogens including endo and ectoparasites such as nematodes, acarians and insects [26]. In comparison with other pediculicides, ivermectin presents some major advantages. First, it has long been recognized that it is a safe pharmaceutical in human health due to low level of toxicity for mammalians, in sharp contrast with classical pediculicides [67]. Indeed, former drugs such as fipronil, lindane or malathion are now forbidden for human use in several countries because of their toxicity and environmental impact [20-25]. Second, ivermectin is currently the only molecule used as an oral treatment against lice. For head lice, oral route avoids suboptimal shampoo application, thus reducing the emergence and transmission of resistant lice [28, 68-70]. For body lice, oral dosing is of particular interest to treat people having a limited access to basic hygiene services. However, despite these remarkable advantages, ivermectin’s efficacy is currently compromised by the emergence of resistant lice [30, 31].

Here, our bioassay confirmed the pediculicidal activity of ivermectin on human body lice. Furthemore, we addressed for the first time the mode of action of ivermectin at the molecular level on glutamate- and GABA-gated receptors. We showed that ivermectin activated only Phh-GluCl channel. In addition, ivermectin blocked the RDL receptor but with a concentration 10 times higher than the concentration required to fully activate Phh-GluCl. These findings suggest that ivermectin do not have any effect on Phh-RDL at the dose used to treat people and reveal that Phh-GluCl is its preferential target. Interestingly, functional silencing using RNAi has already been shown to be an efficient and promising approach to investigate the role of detoxification genes in ivermectin tolerance of human body lice [71]. In order to confirm that GluCl is the main target of ivermectin in lice, it would be interesting to explore *in vivo* the phenotype of *Phh-GluCl* siRNA silenced lice as previously demonstrated for abamectin in *T. cinnabarinus* and in the crop whitefly pest *Bemisia tabaci* [61, 72]. Finally, following the recent observation of GluCl single-nucleotide-polymorphisms (SNPs) in phenotypically ivermectin-resistant head lice [30], our results provide a unique means to explore the molecular mechanisms underlying louse resistance to ivermectin. In a further step, such electrophysiology studies in *Xenopus* oocytes will enable to determine the functional relevance of GluCl SNPs detected in ivermectin-resistant lice [49, 59, 73-75]. Although the putative role of those SNPs are to be determined, we can reasonably assume that mutated subunits can lead to GluCl altered functionality, probably resulting in ivermectin resistance. Additionaly, screening on ivermectin-resistant GluCls recombinantly expressed will provide a powerful tool to optimize the design of new pediculicidal compounds. Further investigations are required to understand the mechanisms of ivermectin resistance and to define new control strategies to avoid emergence of resistance as well as preserve this molecule efficacy.

### On the mode of action of lotilaner in arthropods

With the recent introduction of new generation of compounds in the isoxazoline class (e.g. lotilaner), the development of detailed understanding of the actions of isoxazolines is essential. Arthropod RDL channels are major targets for several old insecticidal classes including dieldrin itself and fipronil [32] and they were recently shown to be the primary target for isoxazolines [39-43]. Therefore, we hypothesized that they could act on RDL from *Pediculus humanus humanus*. In addition to the pharmacological characterization of Phh-RDL and Phh-GluCl, our results show that the inhibitory effects of lotilaner depend on the receptor subtype. Indeed, lotilaner efficiently and selectively blocked the GABA response on Phh-RDL whereas Phh-GluCl was not affected. In order to confirm these findings, it will be interesting to silence the *rdl* gene using RNAi in human body lice [61]. Our results clearly demonstrated the selective antagonistic action of lotilaner on RDL over GluCl channels contrary to A1443 and fluralaner, which were reported to inhibit both RDL and GluCls from *M. domestica* and *R. microplus* [39, 43, 76]. Whether these differences might be due to the compounds themselves or the pharmacological properties of receptors from different species remain to be investigated. Furthermore, the amino acids implicated in the binding of lotilaner are not known yet although some studies recently suggested that amino acids in theTM1 and TM3 might be critical in RDL from *M. domestica* [77]. The mechanism of action of lotilaner on RDL receptors at the molecular level are still to be deciphered. Likewise, further efforts are needed to better understand the molecular basis of the insect selectivity of isoxazolines over vertebrate channels. Contrary to homomeric arthropod RDL channels, the vertebrate GABA are heteromeric channels made of different subunits. Recent studies indicated that A1443, fluralaner and lotilaner were highly selective of arthropod RDL channels while poorly efficient on rat brain membranes (IC_50_ > 10 µM), rat GABA channels (IC_50_ > 30 µM) and canine GABA (IC_50_ > 10 µM), respectively, thus evidencing for low mammalian toxicity [39, 42, 43]. Resistance to isoxazolines has not been reported so far and fipronil resistance is likely not to confer cross-resistance to lotilaner. Likewise, the distinct mechanisms of action for ivermectin and lotilaner seem to indicate that resistance to ivermectin will not extend to lotilaner. Identification of homologous genes encoding RDL subunits and investigation of new expressed recombinant RDL from many other arthropod species including important insect pest species, acari and crustaceans will enhance our understanding of the action of isoxazolines. The potency and selectivity of isoxazolines indicate an important future for this novel class of compounds.

### From dog to human: the future for isoxazoline insecticides as promising pediculicidal candidates for drug repurposing

The isoxazoline class includes a new generation of instecticidal compounds recently introduced for insect chemical control. Since the discovery of isoxazolines, their insecticidal spectrum has become broader to agricultural, sanitary and animal insect pests belonging to eleven orders missing out the order of *Phthiraptera* [39-44]. In the present study, we hypothesized that the isoxazolines could represent attractive candidate drugs to extend the pharmacopeia for louse control. Among the different compounds from the isoxazoline family, we selected the novel insecticide lotilaner as a model for the following reasons: first, it is licensed for protection of companion animals against fleas and ticks and therefore presents a reduced toxicity to mammals; second, it has a long lasting effect after a single oral administration [42, 78]. In dogs, lotiIaner remains active against ticks and fleas during one month with an efficacy ranging from 98 to 100% [79, 80]. Here, we assessed the pediculicidal potential of lotilaner against different stages of body lice. Strikingly, our bioassays on human body lice demonstrated that the adulticide effect of lotilaner was higher than that of ivermectin and fipronil. Whereas 10 µM ivermectin treatment killed 100% of adult lice after 24h, we obtained the same result with 10 µM lotilaner in only 3 hours following exposure. This fast activity is in accordance with the 4h-12h needed to kill ticks and fleas in lotilaner treated dogs [79, 80]. In contrast, neither ivermectin nor lotilaner were found to have a pediculicidal activity on nits. Based on dog data, we can speculate that a single oral dose could provide a pediculicidal activity above 17 days corresponding to the complete life cycle of the human lice [9, 19]. Therefore, the long lasting activity of lolilaner could overcome the absence of activity on embryonated eggs and also prevent re-infestations with a single oral dose. Considering its relatively benign toxicological profile for mammals along with rapid knockdown of lice, lotilaner seems to be an excellent candidate drug to tackle louse infestations in humans. Nevertheless, the innocuity of lotilaner towards human remains to be proven as well as its pharmacokinetics. Interestingly, Miglianico *et al*. recently reported the high efficacy of two other veterinarian drugs from the isoxazoline family (i.e afoxolaner and fluralaner) against a panel of insect species representing major vectors for human tropical diseases [81]. Computational modelling highlighted both the feasibility and the relevancy of using isoxazoline for human medication. In accordance, our results strongly support isoxazoline compounds as major candidates for drug repurposing alone or in co-administration with ivermectin. Indeed, new combinations of active compounds have become frequent in veterinary medicine (e.g. anthelmintics). Alike ivermectin, a useful feature of lotilaner is its mode of administration through oral route, which is more tractable to human than topical formulations. Since ivermectin and lotilaner are efficient at targeting distinct receptors, it is tempting to suggest that a combination of both drugs may have a considerable potential, resulting in an additive or a synergistic effect allowing the control of emerging ivermectin resistant lice and delay or overcome resistance. Such a therapeutic option was recently achieved with the release of fluralaner combined with moxidectin [82, 83] and sarolaner combined with selamectin on the veterinary market for small animals [84, 85]. Hence, it is reasonable to conclude that further investigations on the efficacy of lotilaner alone or in combination with ivermectin for general use as pediculicides would be particularly pertinent and justified. We hope that the results presented here contributes to an increased understanding on the mechanisms behind the physiology of neurotransmission and of the mode of action of the pediculicidal drugs in body lice at the functional level and will provide new therapeutic strategies to ensure the sustainable and effective control of louse infestations.

## Methods

### Chemical assays on body lice

The colony of human body lice (*Pediculus humanus humanus*) were provided by Kosta Y. Mumcuoglu from the Kuvin Center for the Study of Infectious and Tropical Diseases, Hebrew University-Hadassah Medical School, Jerusalem, Israel. Thislaboratory-reared colony adapted to rabbit blood were fed four times per week (permit number of the French Ministry for Research: APAFIS#8455-2017010616224913 v3) and maintained at 30 ± 1 °C and 60-70% relative humidity without exposure to any drugs [86]. Survival and immobilization (knockdown) were monitored at 1, 2, 3 and 24 h after contact of 44 lice with insecticides at concentrations ranging from 1 to 100 µM or 1% DMSO. Nits (50 to 190 per condition) were incubated with lotilaner or ivermectin and hatching was monitored between six and nine days. Water and solution of 1 % DMSO were used as negative controls. Pictures and movies were recorded with a Motic Digital Microscope 143 Series and the Motic Image Plus 3.0 software. Movies were cut out with Apple inc. iMovie 10.1.6.

### Cloning of full-length cDNA sequences of *GluCl* and *rdl* from *Pediculus humanus humanus*

Total RNA was extracted from 30 mg of lice (15 lice) using the Total RNA isolation Nucleospin RNA kit (Macherey-Nagel) according to the manufacturer’s recommendations. Complementary DNA (cDNA) was synthesized using the GeneRacer kit with the SuperScript III reverse transcriptase (Invitrogen) following the manufacturer’s recommendations. Using the *A. mellifera* and *D. melanogaster* GluCl and RDL subunit sequences as querries, tBLASTn searches in NCBI (http://blast.ncbi.nlm.nih.gov/Blast.cgi) allowed the identification of partial genomic contigs (AAZO01006897 and AAZO01005501 for *Phh-GluCl* and AAZO01005738, AAZO01000267 and AAZO01000266 for *Phh-rdl*) and partial mRNA sequences (XM_002429761 for *Phh-GluCl* and XM_002422861.1 for *Phh-rdl*) containing the 3’ cDNA ends. Primers designed on the sequences retrieved from the BLAST searches are reported in **S1 Table**. Then, the corresponding 5’ cDNA ends were obtained by RLM-RACE experiments using the Generacer kit with two rounds of PCR as described elsewhere [87]. After the identification of the cDNA 5’ ends, new primers were designed. The full-length complete coding sequences of *Phh-GluCl* and *Phh-rdl* were subsequently amplified by nested PCRs with the proofreading Phusion High-Fidelity DNA Polymerase (Thermo Scientific) using two pairs of primers. Then, PCR products were cloned into the transcription vector pTB-207 [88] using the In-Fusion HD Cloning kit (Clontech) as described previously [89]. Recombinant plasmid DNA was purified using EZNA Plasmid DNA Mini kit (Omega Bio-Tek) and the sequences were checked (Eurofins Genomics). The novel complete coding sequences of the two Phh-GluCl and Phh-RDL subunits were depositied to Genbank under the accession numbers MT321070 to MT321073. The constructions were linearized with the MlsI restriction enzyme (Thermofisher) and cRNAs were synthetized *in vitro* using the mMessage mMachine T7 transcription kit following the manufacturer’s recommendations (Ambion). Lithium chloride-precipitated cRNAs were resuspended in RNAse-free water and stored at −20 °C.

### Sequence analysis and phylogeny

Signal peptide and transmembrane domain were predicted using SignalP4.1 [90] and SMART [91], respectively. Deduced amino-acid sequences of Phh-GluCl and Phh-RDL were aligned using the MUSCLE algorithm [92] and further processed with GeneDoc (IUBio). The phylogenetic distance trees were generated on amino-acid sequences by the SeaView software [93] using BioNJ Poisson parameters and bootstrap values were calculated with 1000 replicates as described previously [94]. The resulting trees were modified by FigTree (http://tree.bio.ed.ac.uk/software/figtree/). The accession numbers for the protein sequences mentioned in this article are: *Apis mellifera* (Ame): GluCl ABG75737, GRD AJE68942, LCCH3 AJE68943, RDL AJE68941, CG8916 NP_001071290; nAChR7 AJE70265; *Ctenocephalides felis* (Cfe): RDL AHE41088; *Drosophila melanogaster* (Dme): GluCl AAC47266, GRD NP_524131, LCCH3 NP_996469, RDL NP_523991, CG8916 NP_001162770; *Ixodes scapularis* (Isc): GluCl ALF36853; *Musca domestica* (Mdo): GluCl BAD16657, RDL NP_001292048; *Pediculus humanus humanus* (Phh): GluCl1 MT321070, GluCl2 MT321071, RDL1 MT321072, RDL2 MT321073; *Rhipicephalus microplus* (Rmi): GluCl AHE41097, RDL AHE41094; *Tetranychus urticae* (Tur): GluCl BAJ41378 and *Tribolium castaneum* (Tca): GluCl NP_001107775, GRD NP_001107772, LCCH3 NP_001103251, RDL NP_001107809, 8916 NP_001103425.

### Electrophysiology in *Xenopus laevis* oocytes and data analysis

Defolliculated *Xenopus laevis* oocytes were purchased from Ecocyte Bioscience and maintained in incubation solution (100 mM NaCl, 2 mM KCl, 1.8 mM CaCl_2_.2H_2_O, 1 mM MgCl_2_.6H_2_O, 5 mM HEPES, 2.5 mM C_3_H_3_NaO_3_, pH 7.5 supplemented with penicillin 100 U/mL and streptomycin 100 µg/mL) at 19 °C. Each oocyte was microinjected with 57 ng of cRNA encoding Phh-GluCl or Phh-RDL using a Drummond nanoject II microinjector. Three to five days after cRNA injection, two-electrode voltage-clamp recordings were performed with an oocyte clamp OC-725C amplifier (Warner instrument) at a holding potential of −80mV or −60 mV to assess the expression of the GluCl or RDL channels. Currents were recorded and analyzed using the pCLAMP 10.4 package (Molecular Devices). Concentration-response relationships for agonists were carried out by challenging oocytes with 10 s applications of increasing concentrations of compounds. The peak current values were normalized to the response to 1000 µM glutamate or 100 µM GABA giving the maximum current amplitude for GluCl and RDL channels, respectively. For RDL, the effect of antagonists was evaluated, by a first application of each antagonist alone for 10 s, followed by the co-application with 100 µM GABA for 10 s. The observed responses were normalized to the response induced by 100 µM GABA alone performed prior to challenging with the antagonist. The concentration of agonist required to mediate 50% of the maximum response (EC_50_) and the concentration of antagonist required to inhibit 50% of the agonist response (IC_50_) and the Hill coefficient (nH) were determined using non-linear regression on normalized data with GraphPad Prism software. Results are shown as mean +/- SEM and statistical analysis were performed using One-Way ANOVA with Tukey’s Multiple Comparisons Test and paired Student’s t-test. Chloride and sodium permeabilities were conducted as described previously [51]. In brief, the reversal potential of the GABA-induced currents was measured with a 1500-ms-long ramp of voltage from −60 to +60 mV in standard saline recording solution (100 mM NaCl, 2.5 mM KCl, 1 mM CaCl_2_, 5 mM HEPES, pH 7.3) as well as in recording solutions containing a concentration range of either chloride (NaCl replaced by sodium acetate) or sodium (NaCl replaced by tetraethylammonium chloride).

## Materials

Glutamate, GABA, fipronil, picrotoxin and ivermectin were purchased from Sigma-Aldrich. Glutamate and GABA were directly dissolved in recording solution. Fipronil, picrotoxin and ivermectin were first dissolved at 10-100 mM in DMSO and then diluted in recording solution to the final concentrations in which DMSO did not exceed 1 %. Lotilaner (CredelioTM, Elanco, USA) was a commercial formulation purchased in a local pharmacy store. Tablets containing 450 mg lotilaner were dissolved in DMSO to obtain a 100 mM lotilaner stock solution and subsequently diluted in recording solution at a final DMSO concentration less than 0.003%.

## Acknowledgments

We acknowledge the gift of the human body louse colony from Kosta Y. Mumcuoglu from the Department of Microbiology and Molecular Genetics, Kuvin Center for the Study of Infectious and Tropical Diseases, Hebrew University-Hadassah Medical School, Jerusalem, Israel. This study was supported by the Institut National de Recherche pour l’Agriculture, l’Alimentation et l’Environnement (INRAE) and the Université de Tours (annual endowment) and in part by the RTR Fédération de Recherche en Infectiologie (FéRI) of the Région Centre-Val de Loire to CN, CD and IDP. NL is the grateful recipient of a PhD grant from the Animal Health Division of INRAE and from the Région Centre-Val de Loire, France. The funders had no role in study design, data collection and analysis, decision to publish, or preparation of the manuscript.

## Supporting information

**S1 Fig.:**
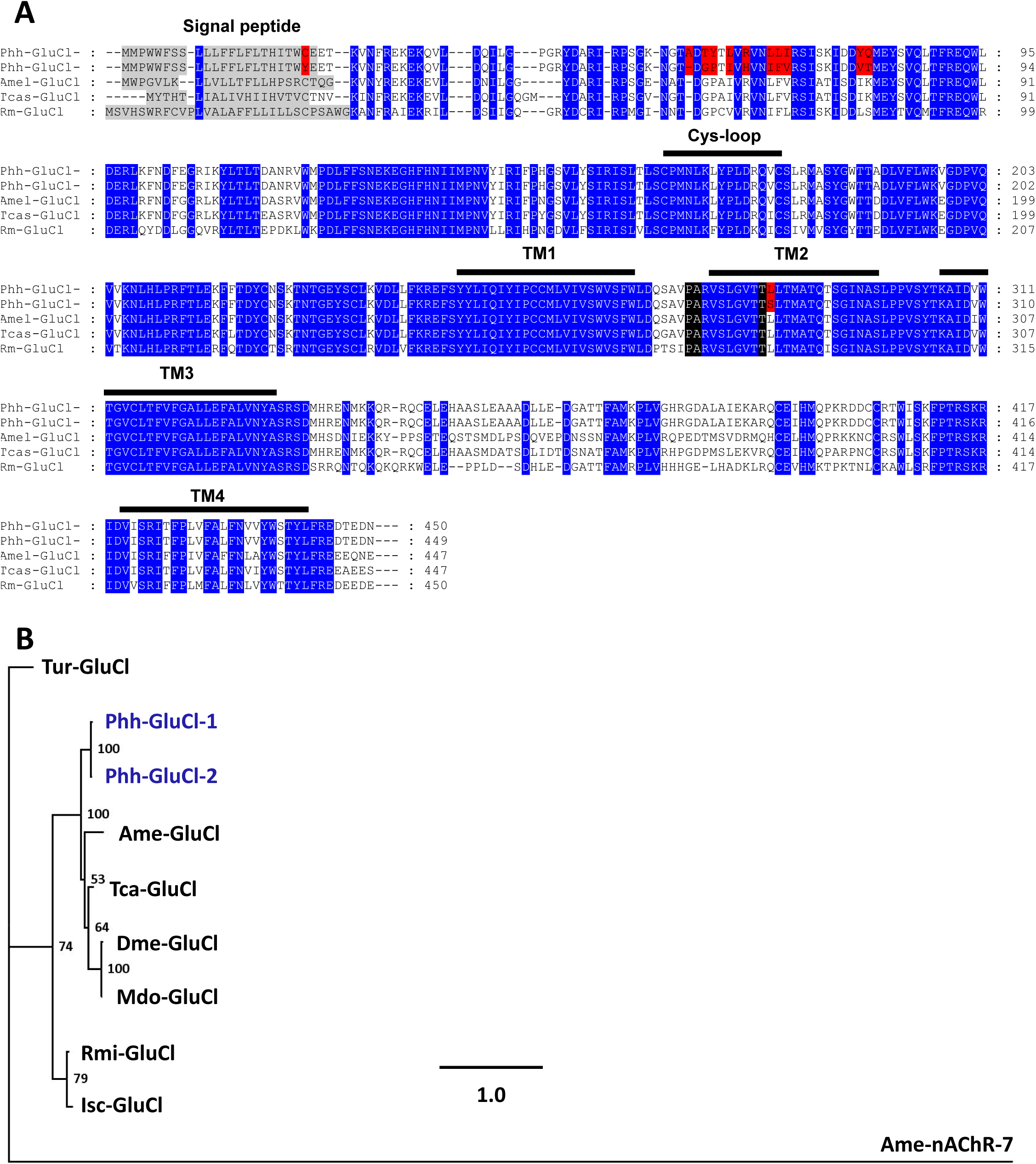
Comparison of insect and acari GluCl subunits. **A**. Alignement of GluCl subunit deduced amino-acid sequences from *Pediculus humanus humanus* (Phh), *Apis mellifera* (Ame), *Rhipicephalus microplus* (Rmi) and *Tribolium castaneum* (Tca). Predicted signal peptides in N-terminal are highlighted in grey. Amino acid differences between the GluCl-1 anf GluCl-2 sequences of *P. humanus humanus* are highlighted in red. Amino acids conserved between all the sequences are highlighted in blue. The cys-loop, transmembrane domains (TM1-TM4) and the highly variable intracellular loop are indicated by the bars. **B**. Distance tree (BioNJ, Poisson) of GluCl protein sequences from insects and acari. The three letter prefixes in gene names Tur, Phh, Ame, Tca, Dme, Mdo, Rmi and Isc refer to the species *Tetranychus urticae, Pediculus humanus humanus, Apis mellifera, Tribolium castaneum, Drosophila melanogaster, Musca domestica, Rhipicephalus microplus* and *Ixodes scapularis*, respectively. The tree was rooted with the *A. mellifera* alpha7 nAChR subunit as an outgroup. Branch lengths are proportional to the number of substitutions per amino acid. Scale bar represents the number of substitutions per site. The bootstrap values are indicated next to each branch. Accession numbers for sequences used in the phylogenetic analysis are provided in the Methods section. The two GluCl sequences of interest are highlighted in blue.

**S2 Fig.:**
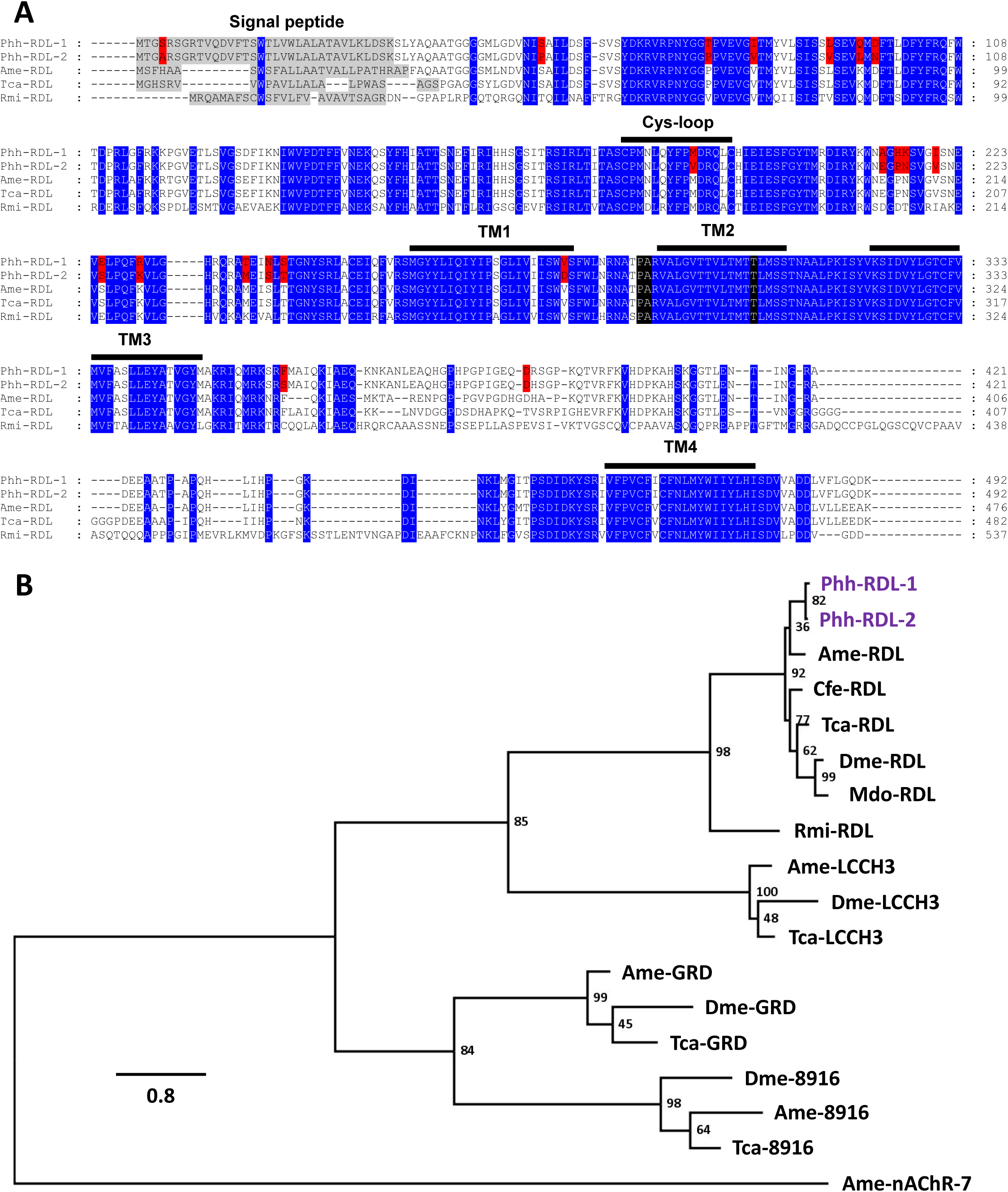
Comparison of insect and acari GABA subunits. **A**. Alignement of RDL subunit deduced amino-acid sequences from *Pediculus humanus humanus* (Phh), *Apis mellifera* (Ame), *Tribolium castaneum* (Tca) and *Rhipicephalus microplus* (Rmi). Predicted signal peptides in N-terminal are highlighted in grey. Amino acid difference between the RDL-1 and RDL-2 sequences of *P. humanus humanus* are highlighted in red. Amino acids conserved between all the sequences are highlighted in blue. The cys-loop, predicted transmembrane domains (TM1-TM4) and the highly variable intracellular loop are indicated by the bars. **B**. Distance tree (BioNJ, Poisson) of GABACl protein sequence from insects and acari. The three letter prefixes in gene names Phh, Ame, Cfe, Tca, Dme, Mdo and Rmi refer to the species *Pediculus humanus humanus, Apis mellifera, Ctenocephalides felis, Tribolium castaneum, Drosophila melanogaster, Musca domestica* and *Rhipicephalus microplus*, respectively. The tree was rooted with the *A. mellifera* alpha7 nAChR subunit as an outgroup. Branch lengths are proportional to the number of substitutions per amino acid. Scale bar represents the number of substitutions per site. The bootstrap values are indicated next to each branch. Accession numbers for sequences used in the phylogenetic analysis are provided in the Methods section. The two RDL sequences of interest are highlighted in purple.

**S3 Fig.:**
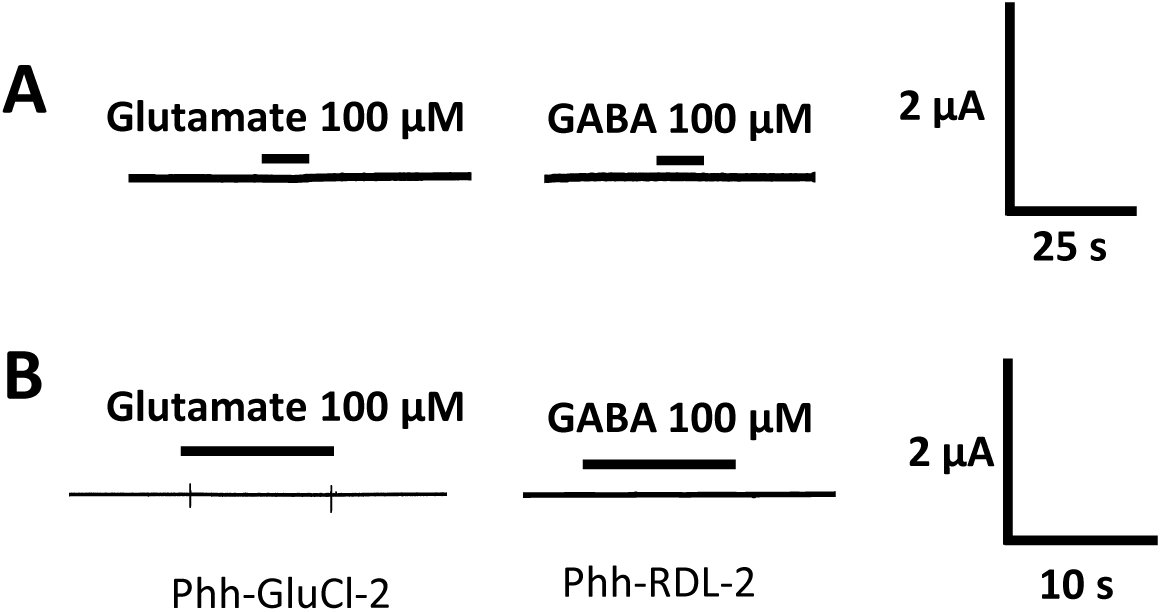
Current traces of uninjected oocytes and oocytes injected with Phh-GluCl-2 or Phh-RDL-2. **A**. Current traces of uninjected oocytes after application of glutamate and GABA at 100 µM. The bar indicates the application time of 10 s. **B**. For Phh-GluCl-2, oocytes were clamped at −80mV and response to 100 µM of glutamate was measured (n = 11). For Phh-RDL-2, oocytes were clamped at −60mV and response to 100 µM of GABA was measured (n = 12). The bar indicates the application time of 10 s.

**S4 Fig.:**
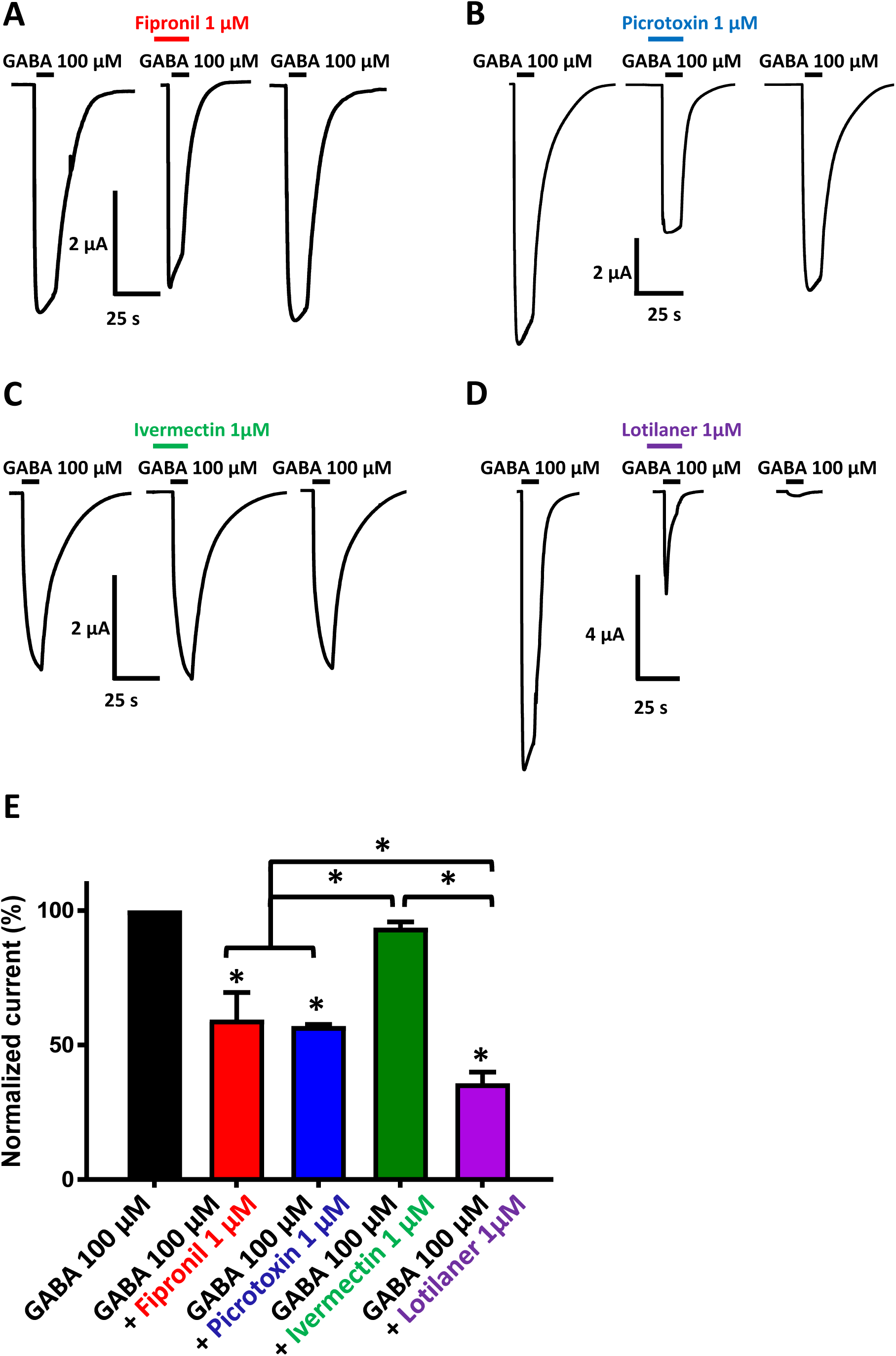
Antagonistic effects of 1 µM of fipronil, picrotoxin, ivermectin and lotilaner on GABA-elicited currents in *X. laevis* oocytes expressing Phh-RDL receptor. **A-D**. Representative current traces evoked by 100 µM GABA with or without co-application of 1 µM fipronil, picrotoxin, ivermectin and lotilaner. Applications of 100 µM GABA alone were performed before and after application of drugs. Drugs were applied for 10 s before the co-application with GABA at 100 µM during 10 s. Application times are indicated by the bars. **E**. Bar chart showing the normalized current responses for 100 µM GABA alone and in the presence of 1 µM fipronil, picrotoxin, ivermectin and lotilaner on Phh-RDL (mean +/- SEM, n = 3-9 oocytes). Currents have been normalized to and compared with 100 µM GABA-elicited currents. * *p* < 0.05; One-way ANOVA.

**Table S1:**
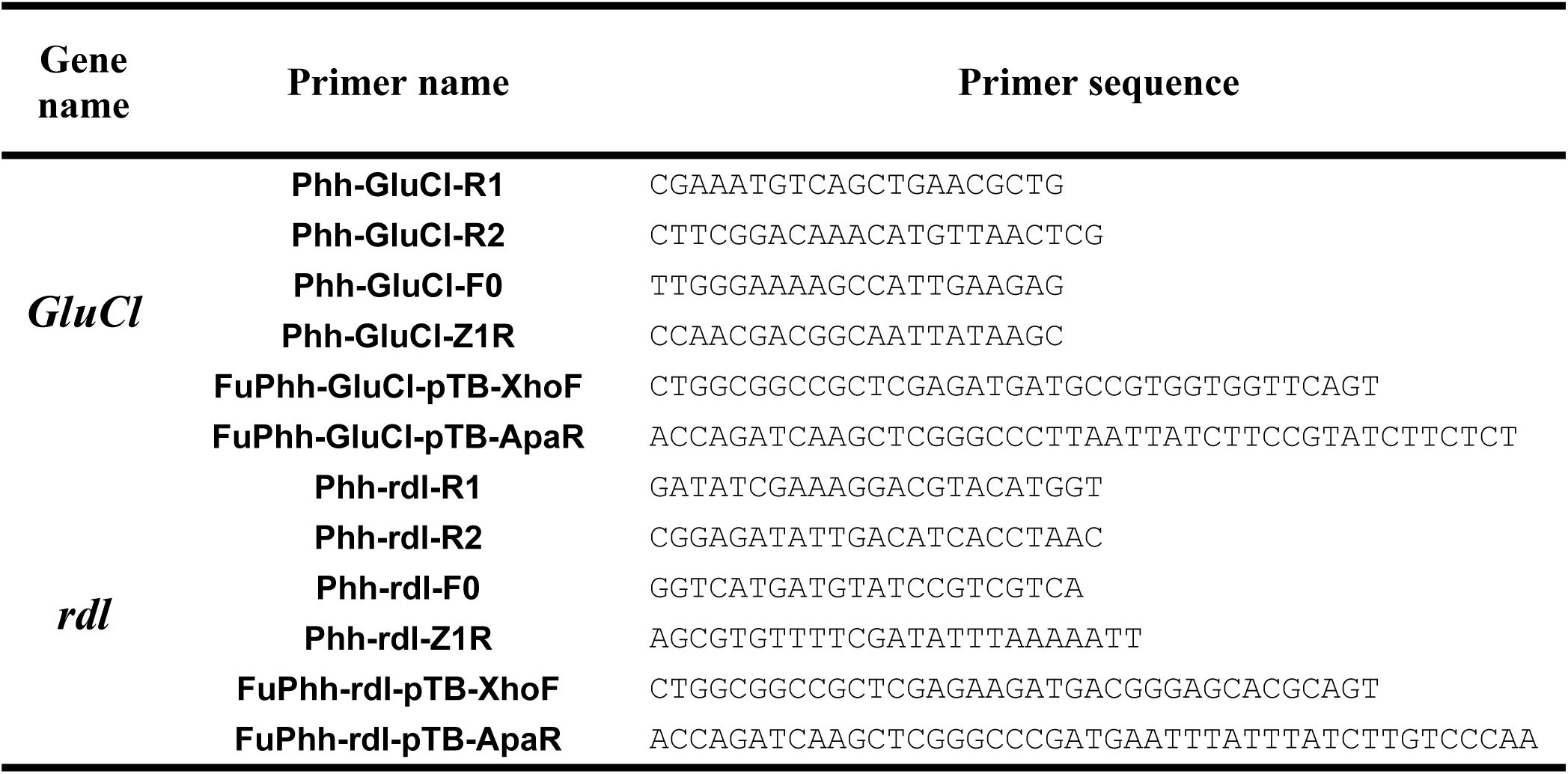
Primers used for the PCR and cloning of *P. humanus humanus* GluCl and RDL subunits

## Author contributions

| Contributor Role | Role Definition |
| --- | --- |
| **Conceptualization** | Ideas; formulation or evolution of overarching research goals and aims.<br>Nicolas Lamassiaude<sup>1</sup>, Berthine Toubate<sup>2</sup>, Cédric Neveu<sup>1</sup>, Françoise Debierre-Grockiego<sup>2</sup>, Claude Charvet<sup>1</sup> Isabelle Dimier-Poisson<sup>2</sup> |
| **Data Curation** | Management activities to annotate (produce metadata), scrub data and maintain research data (including software code, where it is necessary for interpreting the data itself) for initial use and later reuse. |
| **Formal Analysis** | Application of statistical, mathematical, computational, or other formal techniques to analyze or synthesize study data.<br>Nicolas Lamassiaude<sup>1</sup>, Berthine Toubate<sup>2</sup>, Françoise Debierre-Grockiego<sup>2</sup>, Pierre Charnet<sup>3</sup>, Claude Charvet<sup>1</sup> |
| **Funding Acquisition** | Acquisition of the financial support for the project leading to this publication.<br>Cédric Neveu<sup>1</sup>, Berthine Toubate<sup>2</sup>, Isabelle Dimier-Poisson<sup>2</sup> |
| **Investigation** | Conducting a research and investigation process, specifically performing the experiments, or data/evidence collection.<br>Nicolas Lamassiaude<sup>1</sup>, Berthine Toubate<sup>2</sup>, Pierre Charnet<sup>3</sup>, Catherine Dupuy<sup>2</sup>, Claude Charvet<sup>1</sup> |
| **Methodology** | Development or design of methodology; creation of models<br>Nicolas Lamassiaude<sup>1</sup>, Berthine Toubate<sup>2</sup>, Cédric Neveu<sup>1</sup>, Françoise Debierre-Grockiego<sup>2</sup>, Claude Charvet<sup>1</sup> Isabelle Dimier-Poisson<sup>2</sup> |
| **Project Administration** | Management and coordination responsibility for the research activity planning and execution.<br>Claude Charvet<sup>1</sup>, Isabelle Dimier-Poisson<sup>2</sup> |
| **Resources** | Provision of study materials, reagents, materials, patients, laboratory samples, animals, instrumentation, computing resources, or other analysis tools.<br>Berthine Toubate<sup>2</sup>, Cédric Neveu<sup>1</sup>, Claude Charvet<sup>1</sup>, Isabelle Dimier-Poisson<sup>2</sup> |
| **Software** | Programming, software development; designing computer programs; implementation of the computer code and supporting algorithms; testing of existing code components. |
| **Supervision** | Oversight and leadership responsibility for the research activity planning and execution, including mentorship external to the core team.<br>Claude Charvet<sup>1</sup>, Isabelle Dimier-Poisson<sup>2</sup> |
| **Validation** | Verification, whether as a part of the activity or separate, of the overall replication/reproducibility of results/experiments and other research outputs.<br>Nicolas Lamassiaude<sup>1</sup>, Berthine Toubate<sup>2</sup>, Pierre Charnet<sup>3</sup>, Cédric Neveu<sup>1</sup>, Françoise Debierre-Grockiego<sup>2</sup>, Claude Charvet<sup>1</sup>, Isabelle Dimier-Poisson<sup>2</sup> |
| **Visualization** | Preparation, creation and/or presentation of the published work, specifically visualization/data presentation.<br>Nicolas Lamassiaude<sup>1</sup>, Berthine Toubate<sup>2</sup>, Françoise Debierre-Grockiego<sup>2</sup> |
| **Writing – Original Draft Preparation** | Creation and/or presentation of the published work, specifically writing the initial draft (including substantive translation).<br>Nicolas Lamassiaude<sup>1</sup> |
| **Writing – Review & Editing** | Preparation, creation and/or presentation of the published work by those from the original research group, specifically critical review, commentary or revision – including pre- or post-publication stages.<br>Nicolas Lamassiaude<sup>1</sup>, Berthine Toubate, Cédric Neveu<sup>1</sup>, Françoise Debierre-Grockiego<sup>2</sup>, Claude Charvet<sup>1</sup>, Isabelle Dimier-Poisson<sup>2</sup> |

